# Cardiac-sympathetic contractility and neural alpha-band power: cross-modal collaboration during approach-avoidance conflict

**DOI:** 10.1101/2023.10.10.561785

**Authors:** Neil M. Dundon, Alexander Stuber, Tom Bullock, Javier O. Garcia, Viktoriya Babenko, Elizabeth Rizor, Dengxian Yang, Barry Giesbrecht, Scott T. Grafton

## Abstract

As evidence mounts that the cardiac-sympathetic system reacts to challenging cognitive settings, we ask if these responses are passive companions or if they are instead fundamentally intertwined with cognitive function. Healthy human participants performed an approach-avoidance paradigm, trading off monetary reward for painful electric shock, while we recorded simultaneous neural and cardiac signals. Participants were reward-sensitive, but also experienced approach-avoidance “conflict” when the subjective appeal of the reward was near equivalent to the revulsion of the cost. Drift-diffusion model parameters revealed that participants managed conflict in part by integrating larger volumes of evidence into choices (wider decision boundaries). Late alpha-band (neural) dynamics suggested that widening decision boundaries served to combat reward-sensitivity and spread attention more fairly to all dimensions of available information. Independently, wider boundaries were also associated with cardiac “contractility” (an index of sympathetically-mediated positive inotropy). We also saw evidence of conflict-specific collaboration between the neural and cardiac-sympathetic signals. Specific to states of conflict, the alignment (i.e., product) of alpha dynamics and contractility were associated with a further widening of the boundary, independent of either signal’s singular influence. Cross-trial coherence analyses provided additional support for a direct role of cardiac-sympathetics in nurturing fair assessment of information streams during conflict by disrupting the prepotent reward signals. We conclude that cardiac-sympathetic activity is not a mere companion, rather it is a critical component collaborating with cognitive processes to combat reward-sensitivity during the approach-avoidance conflict.

## Introduction

Our nervous system and coupled body evolved together to be flexibly responsive, allowing rapid and often anticipatory changes to meet a broad array of cognitive and physical challenges presented in dynamic environments. Decades of research in cognitive neuroscience have characterized flexible cognitive mechanisms responsible for preserving goal-directed function when external circumstances change. Meanwhile, reactivity in peripheral organ systems, such as the cardiac-sympathetic branch, is well documented in tasks requiring momentary goal-directed changes in mental and physical exertion, showing appropriate reactivity (just enough, just in time) to the task at hand^1–3^. More recent evidence extends cardiac-sympathetic reactivity to complex cognitive challenges such as value-based decision making^4,5^. However it is not yet fully known whether cardiac-sympathetic reactivity is an epiphenomenal companion or whether it is instead fundamentally intertwined with complex cognitive processes. To explore this, we chart the relations between fine-grained behavior parameters, temporally sensitive neural assays and cardiac-sympathetic signals as humans respond to the cognitive challenges presented by an established value-based decision-making paradigm.

We selected the approach-avoidance paradigm^6,7^ for three reasons. First, it creates a challenging state of “conflict”, i.e, when the appeal of the reward is near equivalent to the revulsion of the cost, known to drive lengthier response time (RT) and less consistent choice behavior^4,8,9^ (Figures 1A-B). Second, the paradigm can identify prepotent sensitivity toward a particular value dimension, such as the reward sensitivity commonly reported in healthy controls^4,9–11^ (Figure 1C). Third, conflict arising from integrated reward and cost recruits a broad array of frontal cortical brain networks^4,8,9,12–16^, in addition to increased cardiac-sympathetic responses^4,5^, suggesting the paradigm drives qualitative shifts both in behavior and across a complex physiological cascade, allowing us to test cross-modal relations. To decompose behavior even further, and extract fine-grained assays of behavioral perturbations to correlate with physiology signals, we fitted the drift-diffusion model^17–25^ (DDM) to choice and RT data (Figure 1D). Initially considered in perceptual contexts^17–19^, parameters from the DDM are an increasingly useful tool for disambiguating the underlying reasons for lengthier RT in more complex value-based contexts^20–25^ (such as people incorporating more information into decisions vs accumulating information less efficiently - Figure 1D).

**Figure 1.**
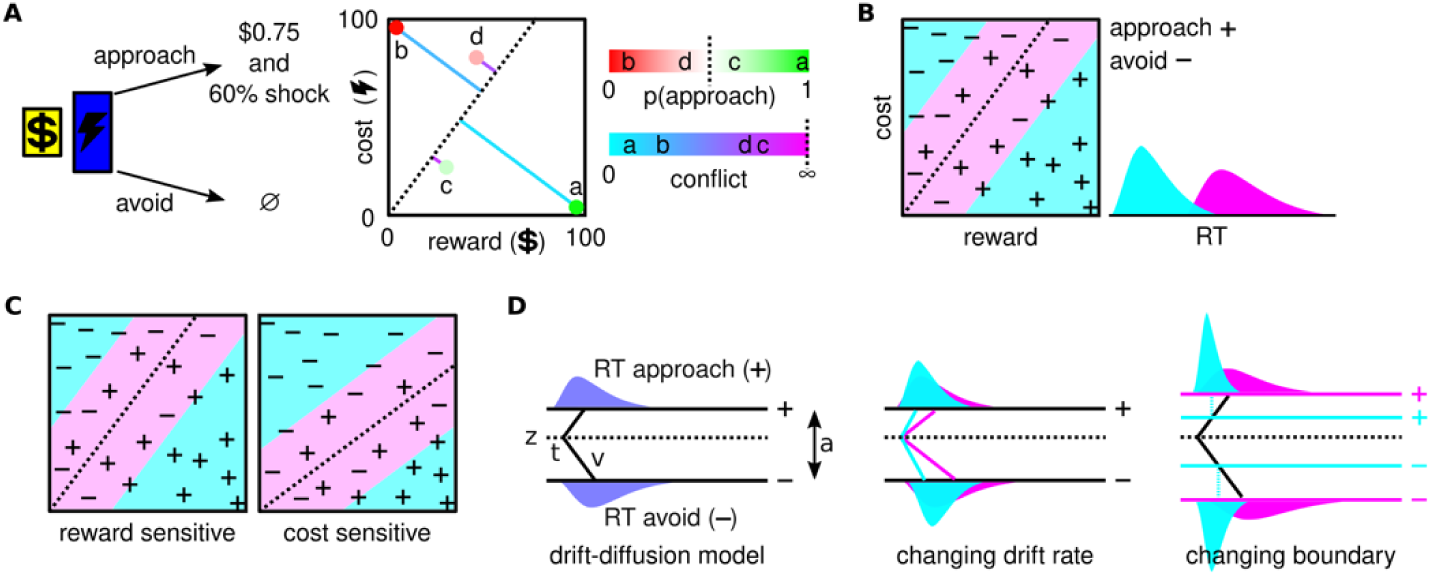
Approach-avoidance and drift-diffusion model frameworks. (A) In the approach-avoidance paradigm participants integrate a reward and a cost in a “take-both-or-leave-both” choice regarding a compound offer. Varying the levels of reward and cost over multiple offers affords a two-dimensional logistic framework that can identify subjective value (p(approach); red-green gradient) and “conflict” (aqua-fuchsia gradient) across the decision space. Conflict is maximal near a “threshold” (dashed line), i.e., as p(approach) nears 0.50. Four example offers are shown (a-d) that vary in subjective value and conflict. (B) High conflict (fuchsia) typically makes choices less consistent with lengthier RT. (C) The slope of the “threshold” characterizes a sensitivity for reward or cost. Fitting the logistic model separately for each participant accounts for such sensitivities prior to enumerating where in decision space they subjectively experience conflict. (D) The drift-diffusion model assumes choice and RT data can be modeled as a sequential sampling process; following an initial nondecision time (t), the decision process begins at starting point (z) and accumulates evidence at rate (v) toward one of two boundaries that determines the choice (in our case, approach (+) or avoid (-)); boundaries are separated by a distance (a). Parameters provide a fine-grained assay of behavior, such as any bias toward one choice (z), how rapidly evidence is integrated during decision formation (v) or the amount of evidence required before a choice is executed (a wider boundary denoting a more conservative criterion). States of high conflict might impact any or all of these parameters. We depict simulated schematics (n=1000 trials) of singularly changing the drift rate or the boundary separation. In each, we fixed a set of baseline parameters (t = 0.30; v=1; a=2; z=0.60), and then increased or decreased v or a by 40%. Note that in each panel, there is a bias toward approach (z > 0.50), and identifiably different features in the RT distributions of approach and avoid resulting from the parametric changes. For more in-depth examples, see^18^.

We used a graded version of the paradigm^26^ to measure neural responses during early and late stages of decision formation. We also applied selective frequency “tagging” and lateralized positioning of the reward and cost information to extract steady-state visually evoked potentials (SS)^27–31^ and spatially responsive dynamics in the alpha band^32–34^. For cardiac-sympathetic variables, we recorded beat-by-beat estimates of contractility (inotropy), which is primarily mediated by noradrenergic sympathetic-autonomic drive and associated with reactivity to challenge^35–41^. Together, these neural and cardiac-sympathetic variables form a broad profile of physiological responses and can capture any interplay involving sensory-gain (SS), goal-directed attention (alpha), and more peripheral sympathetic responses (contractility).

Consistent with previous work, our sample of healthy human participants were primarily reward-sensitive, but showed evidence of altered behavior when encountering high approach-avoidance conflict. They appeared to computationally respond to this high conflict by requiring more evidence before committing to choices (wider decision boundary), reducing a bias toward approach (lower starting point) and integrating evidence more slowly (slower drift rate). Behavioral features could further be predicted trial-by-trial by both neural (primarily late alpha) and cardiac-sympathetic responses, with the majority of features related to changes of the decision boundary. Regardless of state (i.e., low or high conflict), wider boundaries, indicating a pursuit of greater evidence for choices, were first associated with late alpha-band dynamics consistent with attentional processes combating reward-sensitivity and spreading attention more fairly to all dimensions of available information. Independent of the alpha relations, wider decision boundaries were also associated with increased cardiac contractility, suggesting “cross-modal collaboration”, i.e., that neural and cardiac-sympathetic systems were independently recruited by the pursuit of evidence. Critically, and unique to states of high conflict, we observed interactive cross-modal collaboration, that is, “alignment” between alpha-band dynamics and contractility modulating the boundary above the singular influence of the two systems in parallel. In a separate analysis using raw data (cross-trial alpha-band coherence), we corroborated that the interaction of alpha and contractility is relevant primarily in high conflict. We additionally reveal that their interaction might serve to disrupt dynamics in the prepotent information signal (reward), thus characterizing a potential (and more directly mechanistic) role of sympathetic activity in nurturing fair assessment of information streams during conflict. We conclude that cardiac sympathetic activity is unlikely a mere companion. Instead, it may be a critical component operating alongside cognitive processes to combat reward sensitivity during the approach-avoidance conflict.

## Results

We recorded continuous multi-channel electroencephalography and cardiac-sympathetic physiology (combined electrocardiography and impedance cardiography) while human participants made approach-avoidance choices regarding offers that varied trial-by-trial in reward and in cost. Each “take-it-or-leave-it” trial offer gradually presented a monetary reward ranging in value from $0.01 to $1.50 and a shock cost ranging in value from minimal to near maximum bearable pain (see trial schematic in Figure 2A). On average, participants accepted 68% (σ=12.6%) of offers, and responded with median RT of 1.73 seconds (σ =0.428) relative to offer onset.

**Figure 2.**
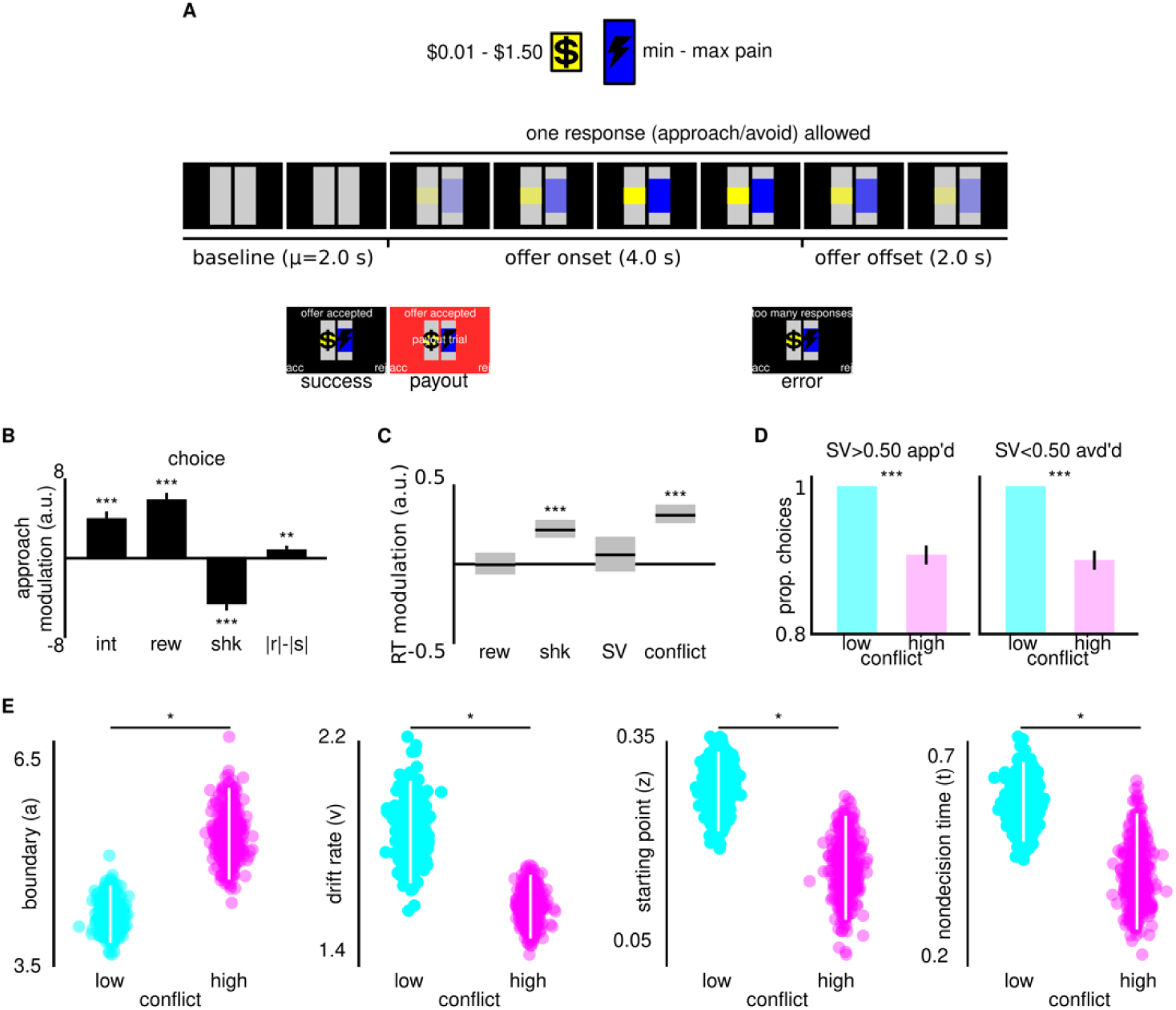
Graded approach-avoidance paradigm reveals fine-grained behavioral responses to conflict. (A) Participants approached (accept) or avoided (reject) offers pairing varying levels of monetary reward with varying levels of painful electric shock (communicated via size of relevant bar) with a single response during gradual onset of stimuli; see Methods for success, payout and error trials. (B) Participants integrated reward (rew) and cost (shk) into choices, with a greater weighting of reward (|rew|-|shk| > 0), indicating reward sensitivity. Error bars are standard error of the mean across parameter estimates for each subject. ***p<0.001, ***p<0.01. (C) Response time (RT) was longer when offers offered higher cost (shk) or higher conflict, but not higher objective (rew) nor subjective (SV) reward. See Methods and Figure 1A for how SV (i..e, p(approach)) and conflict are enumerated trial-by-trial. Thick vertical grey lines span highest density interval (HDI) of coefficient posterior. Thin black line is posterior median. ***HDI of coefficient posterior does not contain 0. (D) Responses were more inconsistent in high vs. low conflict. Lower proportion of choices with SV>0.50 approached (app’d; left) and with SV<0.50 avoided (avd’d; right). Error bars are standard error of the mean across subjects. ***p<0.001. (E) In states of high conflict, participants sought more evidence (boundary - (a)) had a lower rate of evidence accumulation (v), had less of a bias toward approach (starting point (z)) and had slightly shorter nondecision time (t). Boundary units are arbitrary “evidence”, and drift rate is in units of “evidence” per second; starting point (z) is on a logit scale where positive values (i.e., >0.50) are closer to approach boundary (see caption for Figure 1D). nondecision time (t) is measured in seconds. Digitized violin plots contain 400 samples from parameter posterior. Vertical white lines span posterior HDI. *credible Bayesian difference between two parameters (θ1,θ2), i.e., 0∉HDI(D(θ1,θ2)), where D=P1(θ1∣X1)−P2(θ2∣X2).

### Behavioral results

Choice and response-time (RT) analyses confirmed that participants were reward sensitive and also confronted approach-avoidance “conflict” performing our task. From two-dimensional logistic models fitted separately to each individual subject’s set of choices, modeling p(approach) as a function of an intercept (b_0_) and the magnitude of monetary reward (b_1_) and shock cost (b_2_) offered on each trial, these two continuous value dimensions respectively parametrically increased (group-mean b_1_=6.05; t(26)=9.55; p<0.001, Figure 2B) and decreased (group-mean b_2_=-4.76; t(26)=-7.65; p<0.001, Figure 2B) the log odds of approach. In other words, participants integrated both a reward and a cost into their choices. However these parameters also confirmed that participants were reward sensitive, characterized by a bias toward approach (group-mean b_0_=4.15; t(26)=6.08; p<0.001, Figure 2B), and an overweighting of reward in the integration of value dimensions (group-mean |b_1_|-|b_2_|=0.987; t(26)=3.59; p<0.001, Figure 2B), consistent with previous studies^19,20^.

We next confirmed the typical behavioral features of encountering conflict - lengthier RT and inconsistent choice. First, the best fitting model of participants’ log-transformed trial-by-trial RT contained a credibly positive group-level coefficient for conflict (lE(β)=0.304; HDI(β)=[0.245,0.363]; Figure 2C). RT was therefore credibly longer on offers that were closer to the decision threshold depicted in Figure 1A, i.e., presenting a higher level of conflict. We also observed a credibly positive group-level coefficient for cost (lE(β)=0.215; HDI(β)=[0.159,0.270]; Figure 2C), offering additional evidence that participants were reward-sensitive at the group level, i.e., lengthier deliberation with increased cost. RT was not credibly modulated by either the objective nor subjective level of reward on offer (Figure 2C). We next confirmed the second typical behavioral feature of conflict - increased choice inconsistency. This was confirmed at the group level, via both lower acceptance of positive subjective value (t(26)=7.2038; p<0.001) and lower avoidance of negative subjective value (t(26)=6.97; p<0.001) in states of high conflict (Figure 2D).

We next fitted a baseline hierarchical Bayesian^42^ drift-diffusion model (DDM) to get a clearer insight into how increased conflict alters the computational parameters governing choice and RT. This model fitted distinct group-level DDM parameters [a,v,z,t], depending on whether participants were making choices on trials binned into states of low or high conflict. This baseline model revealed that in high conflict, participants sought more evidence before executing their choices (a wider decision boundary (a), Figure 2E), and additionally reached this decision boundary by way of a dampened rate of evidence accumulation ((v) - Figure 2E). Starting points (z) in this DDM also confirmed an overall bias toward approach in states of both low and high conflict, however this bias was attenuated in high conflict (Figure 2E). Finally, nondecision time (t) was slightly shorter in high conflict (Figure 2E).

Summarizing the behavioral results from the approach-avoidance paradigm, participants were reward sensitive, but also confronted states of subjective “conflict”. A baseline DDM additionally suggested that the parametric-behavioral response to states of high conflict involved a larger requirement of evidence prior to committing choices, a slower accumulation of evidence toward that criterion and an attenuated bias toward approach behavior.

### Cross-modal (neural and cardiac-sympathetic) collaborative influence of drift-diffusion mode (DDM) parameters

The above baseline model summarized group-level DDM parameters [a,v,z,t] over all trials in states of either low or high conflict, however parameters governing behavior are dynamic and can vary on a moment-to-moment basis^42^. We therefore tested if trial-by-trial perturbations in DDM parameters could be predicted by trial-by-trial physiological responses (Figure 3A-C). First, by flickering the reward and cost information at specific frequencies, we could extract timeseries for steady-state visually evoked potentials relevant for reward (SS_rew_) and cost (SS_shk_) information, in addition to a “symmetry” trace (SS_sym_) enumerating the similarity between these traces (i.e., the symmetry across the brain) over time (Figure 3A). Next, and by also lateralizing the reward and cost stimuli, we extracted timeseries of spatially-sensitive alpha power, i.e., relevant for reward (alpha_rew_) and cost (alpha_shk_) information, in addition to a “symmetry” trace (alpha_sym_) enumerating their similarity over time (Figure 3B). For each SS and alpha variable, we computed the average power in early ([0 s to 1 s] post offer onset) and late ([1 s to 2 s]) time windows, making a total of 12 neural variables for each trial (Figure 3A-B). Finally, by recording simultaneous impedance cardiography (ICG) and electrocardiography (EKG), we computed an estimate of cardiac contractility (inotropy), i.e., the time window between initial systolic electrical innervation of the heart and the mechanical opening of the aortic valve on each heartbeat (pre-ejection period “PEP” Figure 3C). We averaged across a positively-scored contractility estimate for each beat registered in the two-second window post offer onset (Figure 3C).

**Figure 3.**
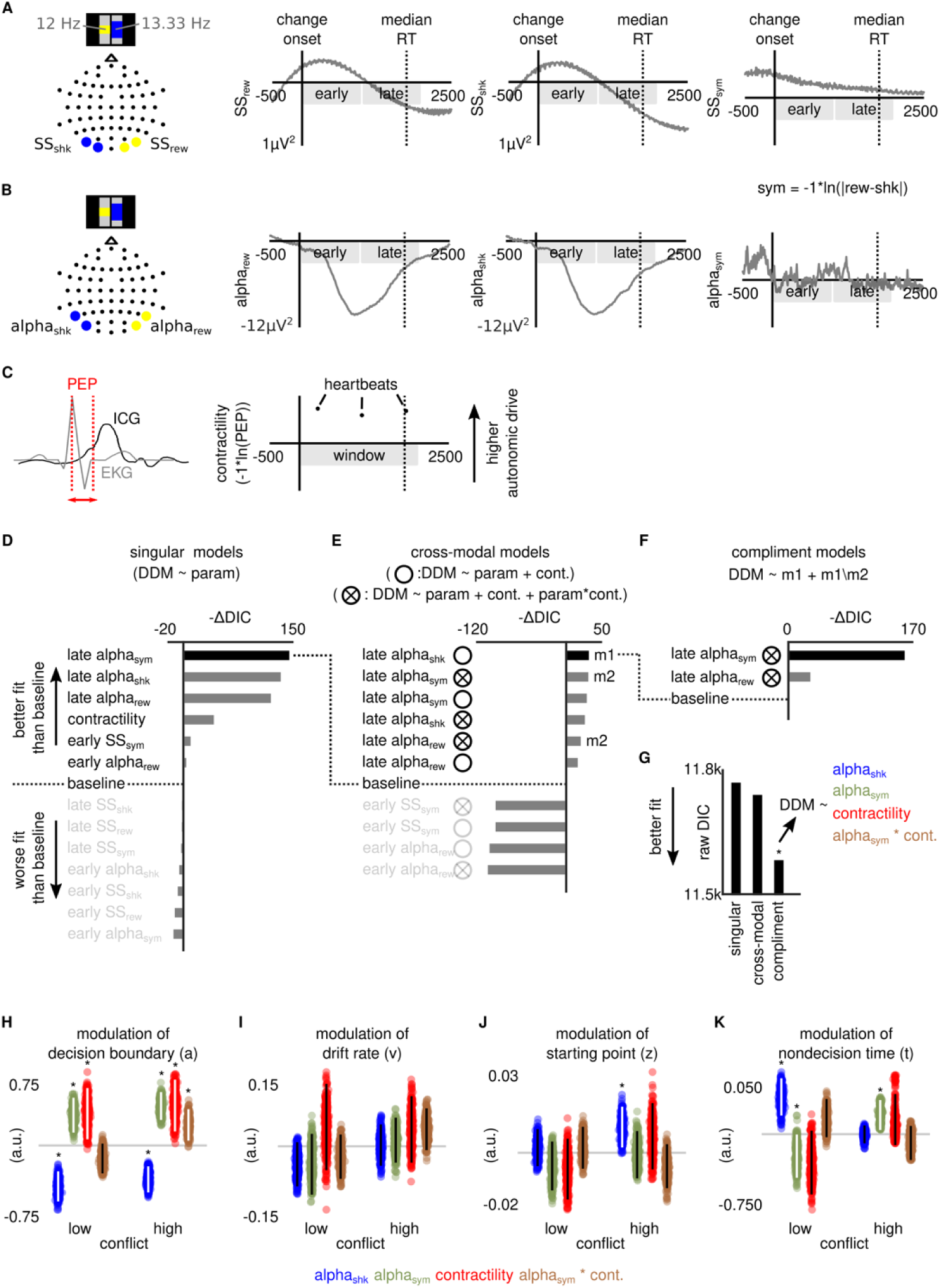
Interactive cross-modal collaboration influences the decision boundary of the drift-diffusion model (DDM) (A) Separate flicker-rates applied to reward and cost stimuli afforded capture of steady-state visually-evoked potential timeseries for reward (SS_rew_) and cost (SS_cst_). In the “symmetry” timeseries (SS_sym_), higher values reflect greater symmetry (more equal power) between the two SS timeseries (-1*ln|(SS_rew_-SS_shk_|)). Timeseries were averaged in early [0 to 1 s] and late [1 s to 2 s] time windows relative to offer onset. (B) Lateralized stimuli afforded capture of alpha-power timeseries relevant for reward (alpha_rew_) and cost (alpha_cst_). In the “symmetry” timeseries (alpha_sym_), higher values reflect greater symmetry (more equal power) between the two alpha timeseries (-1*ln|(alpha_rew_-alpha_shk_|)). Timeseries were averaged in early [0 to 1 s] and late [1 s to 2 s] time windows relative to offer onset. (C) The pre-ejection period (PEP) is recorded with combined impedance cardiography (ICG) and electrocardiography (EKG); shorter PEP indicates increased sympathetic beta-adrenergic myocardial contractility. Our contractility estimates, where higher values reflect greater cardiac-sympathetic drive (contractility=-1*ln(PEP)), were averaged across each heartbeat in a [0 to 2 s] time window relative to offer onset. (D) Singular models predicted each DDM parameter [a,v,z,t] by the state of either a neural variable or contractility on each trial, separately for states of low and high conflict (eight regressors in total). Six models improved fits beyond the baseline model in Figure 2E. Fits assessed relative to baseline with improvements in deviance information criterion (-ΔDIC), positive values reflecting better fit. (E) Cross-modal models predicted each DDM parameter [a,v,z,t] with either additive or interactive models winnowed from the fits in Figure 3D. Additive models (empty circles) predicted DDM parameters by the state of a neural variable in addition to contractility (separately for states of low and high conflict; sixteen regressors in total), while interactive models (circles with crosses) also included a third regressor for the product of the neural signal and contractility (24 regressors in total). Six models improved fits beyond the best fitting model in Figure 3D. Fits assessed relative to baseline with improvements in deviance information criterion (-ΔDIC), positive values reflecting better fit. (F) Compliment models asked if the fit of the best fitting cross-modal model (which included alpha_shk_; marked “m1” in Figure 3E) could be improved by adding the compliment (i.e., set difference) from either models including alpha_rew_ or alpha_sym_ (marked “m2” in Figure 3E). Both improved the fit beyond the best fitting model in Figure 3E. Fits assessed relative to baseline with improvements in deviance information criterion (-ΔDIC), positive values reflecting better fit. (G) The best overall fitting model was the compliment model from Figure 3F, where DDM parameters were predicted by four regressors: alpha_shk_, contractility, alpha_sym_ and alpha_sym_*contractility. Fits assessed with raw deviance information criterion (DIC), lower values reflecting better fit. (H-K) Parameter posteriors from best fitting (compliment) model of DDM parameters. Most neural and cardiac-sympathetic relations involved the decision boundary (a). In low and high conflict, relations suggested wider boundaries indicated a sympathetic-supported fairer spread of attention to all dimensions of available information; greater desynchronization of alpha_shk_, greater symmetry in alpha (alpha_sym_) and increased contractility. Unique to states of high conflict, the boundary showed additional positive modulations with interacting cross-modal signals (alpha_sym_*contractility(cont.)). Digitized violin plots contain 400 samples from parameter posterior. Vertical lines span highest density interval (HDI) of coefficient posterior, and are white if HDI does not contain 0 (also marked with *), black otherwise.

We used an iterative modeling approach to find a best fitting combination of physiological signals predicting DDM parameters on each trial, with an emphasis on discovering cross-modal (i.e., neural and cardiac) collaboration (Figure 3E-H). In a series of singular models (Figure 3E) we first asked if DDM parameters could be predicted by a single regressor. Each singular model tested the relation between one of the above 13 physiological variables and each of the four DDM parameters [a,v,z,t], allowing for a separate relation for trials in states of low and high conflict (i.e., each of the 13 singular models tested eight relations in total). Using Deviance Information Criterion scores (DIC^42^), we determined if any of these singular models offered an improved fit of choice and RT data relative to the baseline model fitted in “Behavioral results” above (i.e., fitting static estimates of a,v,z and t in each conflict state). From resulting differences in DIC scores, and as depicted in Figure 3D, six models provided an improved fit; these model respectively predicted trial-by-trial DDM parameters with late alpha_sym_, late alpha_shk_, late alpha_rew_, contractility, early SS_sym_, and early alpha_rew._ We therefore established that momentary perturbations in behavior, as decomposed by the DDM, could be predicted by both neural and cardiac-sympathetic physiological signals, with neural signals predominantly involving the alpha band, and signals measured in the later time window.

The above six singular models reveal a better fit of data predicting DDM parameters by either a neural or cardiac-sympathetic variable. To test cross-modal collaboration, we next asked if any of the 5 singular (i.e., single-regressor) neural models could be improved by adding contractility as a second regressor (i.e., cross-modal vs. singular model). We also tested for evidence of both additive and interactive cross-modal collaboration. For additive collaboration, we tested models with two regressors (the neural variable in question, and contractility), while for interactive collaboration, we added a third interaction or ‘alignment’ regressor which was the normalized product of the neural variable in question and contractility. This made ten target cross-modal models in total (Figure 3E). We also updated the baseline, and tested if these cross-modal models improved the fit relative to the best fitting singular model, i.e., which had alpha_sym_ as its sole regressor (dashed line from Figure 3D to 3E). From resulting differences in DIC scores, we observed six cross-modal models providing an improved fit over baseline. All such models involved alpha-band activity recorded in the late window (Figure 3E). The best fitting model was additive; it contained alpha_shk_, that is, alpha power relevant for the cost information, alongside contractility, and no third alignment regressor.

The set of cross-modal models revealed that behavioral features could be better predicted by a neural variable relative to cost (alpha_shk_) in addition to cardiac-sympathetic perturbations (contractility), i.e. side-by-side in the same model. However, given that cross-modal models involving alpha power relevant to reward information (alpha_rew_) and the symmetry of alpha across the brain (alpha_sym_) also provided an improved fit relative to baseline (Figure 3E), we ran a final set of models to test for their complimentary modulation of DDM parameters. In other words, we tested if the outright best cross-modal fit (additive alpha_shk_, marked “m1” in Figure 3E) could be improved even further by also including compliment (i.e., set difference) parameters from either of the two best-fitting cross-modal models involving alpha_rew_ and alpha_sym_ (each of which was interactive; each marked “m2” in Figure 3E). We again updated the baseline to the best-fitting cross-modal model (dashed line from Figure 3E to 3F). From resulting differences in DIC scores (Figure 3F), we observed both complimentary models to improve the fit, with substantial improvement in the case of adding set difference parameters involving the interactive cross-modal model with alpha_sym._. In other words, the best fitting compliment model allowed DDM parameters to be predicted not just by alpha_shk_ and contractility, but also by the symmetry of alpha across the brain (alpha_sym_) and the product of alpha_sym_ and contractility (Figure 3F-G). Our iterative modeling approach therefore unearthed a set of both complex neural (albeit exclusively alpha) and cardiac-sympathetic physiological signals that predicted parameters of the DDM, and further revealed evidence for interactive cross-modal collaboration (i.e., neural and cardiac-sympathetic alignment).

Inspecting the parameter posteriors of this best-fitting compliment model (Figure 3H-K), the majority of modulations occurred with the decision boundary (a). Coefficients with neural signals were consistent with wider boundaries accompanying a fairer spread of attention to all dimensions of available information, though this wasn’t exclusive to states of high conflict. In both low and high conflict states, there was a negative association between the boundary and alpha relevant to cost (alpha_shk_; Figure 3H), suggesting greater desynchronization of alpha contralateral to cost information as the boundary increased (consistent with more attention being allocated to that information). Also in both states, there were positive associations between the boundary and the symmetry of alpha on either side of the brain (alpha_sym_; Figure 3H). Widening boundaries were therefore not solely associated with deploying attention to cost, but also driving a more even spread of attention to both channels of information, consistent with a pursuit of greater evidence, but in an additive, i.e., not over-riding manner. The relationship between the boundary and cardiac contractility was likewise observed in both states, with this (positive) association consistent with wider boundaries also accompanying increased sympathetic drive (contractility; Figure 3H). Exclusive to states of high conflict, however, we saw the interactive aspect of cross-modal collaboration, i.e., neural and cardiac-sympathetic signals aligning with meaningful impact on behavior. That is, in addition to its singular relations with alpha_sym_ and contractility, the boundary was additionally modulated positively by their alignment. Strikingly, this interaction only occurred in states of high conflict (alpha_sym_*cont.; Figure 3H). In other words, as participants pursued greater evidence to execute choices near their decision threshold (Figure 1A), there was a separate likelihood of increased cross-modal collaboration, i.e., greater phasic alignment of the cardiac-sympathetic (positive inotropic) response and the degree of alpha symmetry across the brain. This was the sole evidence of such interactive cross-modal collaboration in our best-fitting model.

We observed more sparse influence of the physiological signals across the remaining DDM parameters. Drift rate (v) was not credibly modulated by any signal (Figure 3I). The starting point (z) was modulated only in states of high conflict by alpha related to cost (alpha_shk_; Figure 3J). This positive association is consistent with a bias toward approach intensifying when less attention is directed at the cost information (i.e., more synchronization of alpha_shk_). Finally, state-specific modulations emerged relating to nondecision time (t). Nondecision time is a constant term included in the DDM to account for early perceptual and motor preparation processes. In low conflict, nondecision time was lengthened by both a decrease in alpha symmetry, and more synchronization of alpha relevant to cost (alpha_shk_ and alpha_sym_; Figure 3K). Conversely, in high conflict, nondecision time was lengthened solely by the increase in alpha_sym_ (Figure 3K).

### Raw-data corroboration of cross-modal interaction using inter-trial alpha-phase coherence

In the above section we observed that power in the alpha band operates both side-by-side and in interaction with cardiac-sympathetic signals, consistent with them supporting an attentional response to both a reward sensitivity (i.e., deploying more attention to cost) and more generally resolving a broader conflict (spreading attention more fairly to all information). We next sought corroborative evidence of such cross-modal resolution of conflict using a raw form of data independent from computational models. Additionally, we sought additional insights into putative functional mechanisms. With this in mind, we probed how the state of cardiac-sympathetics (specifically, whether contractility was high or low) impacted the coherence of alpha power across trials, relevant to both the reward and cost information. We re-processed the alpha-band activity contralateral to the reward and cost information, extracting the phase angle of these waveforms over time, i.e., alpha_rew_θ and alpha_shk_θ. If specific events (such as the onset of an offer) evoked temporally consistent fluctuations in alpha power, phase angles would summarize across trials to a coherent sinusoid-like waveform oscillating in and around the alpha-band frequency (7Hz - 14Hz). Figure 4A depicts alpha_rew_θ and alpha_shk_θ summarized across trials and participants, separately for trials that were above (high contractility) or below (low contractility) a participant’s median for trials in states of high conflict. Here we observe alpha-like oscillations present in each time series. However during the later time window, th sinusoidal patterning in alpha_rew_θ (Figure 4A top panel) appears to be greater on low-contractility trials. A three-way within-subjects ANOVA of summarized rectified phase angles |θ| as a function of the timeseries (alpha_rew_θ, alpha_shk_θ), time window (early, late) and contractility (high, low) confirmed by way of a three-way interaction (F(1,26)=5.84, p=0.023) that coherence wa indeed higher in low-contractility (|θ|=0.117) vs high-contractility (|θ|=0.091; p_Tukey_=0.012), only in the alpha_rew_θ timeseries and only in the late time window (Figure 4B). In addition, the same three-way ANOVA returned no three-way interaction for trials in states of low conflict (F(1,26)=0.5033, p=0.484). Thus, in this corroboration analysis using both raw data and an alternative means to look at frequency decomposition (coherence vs power), we first confirmed that the interaction of alpha and contractility is relevant primarily in states of high conflict. We additionally reveal a potential mechanistic role of cardiac-sympathetics in the cross-modal collaboration that drives a fairer appraisal of information: disrupting dynamics in the prepotent information signal of reward sensitivity.

**Figure 4.**
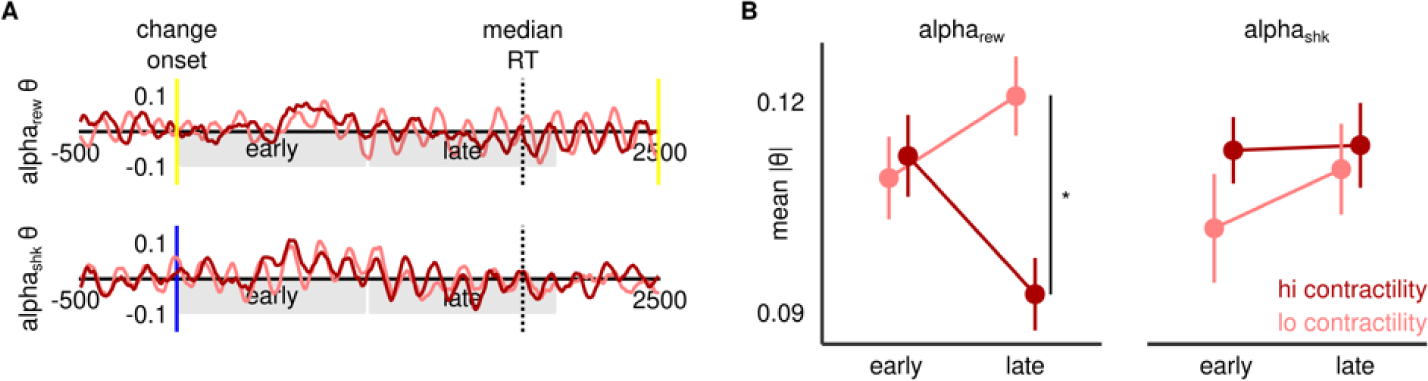
Reward-related alpha-band coherence selectively attenuated by contractility in high conflict. (A) Phase-angle timeseries of alpha contralateral to reward (alpha_rew_θ - top) and cost (alpha_shk_θ - bottom) in high conflict, averaged across subjects separately for trials that were higher (dark red) or lower (light red) than their median contractility. (B) Summarizing phase coherence (absolute phase-angle value |θ|) across early and late time-windows, we see a three-way interaction whereby late coherence diminishes significantly in high contractility, and only in the alpha timeseries contralateral to reward.

## Discussion

Event-related physiological sciences have laid the foundations to explore cross-modal (i.e., neural and cardiac-sympathetic) collaboration subserving complex value-based behavior. We recorded parallel continuous electroencephalographic and cardiac-sympathetic data to probe an interplay involving sensory-gain (SS), goal-directed attention (alpha), and cardiac-sympathetic responses (contractility) while humans performed a modified version of the approach-avoidance paradigm. Participants were reward sensitive but encountered “conflict” when approach and avoidance presented similar value. Using the drift-diffusion model (DDM), we computationally decomposed their behavioral adjustments to conflict, which principally involved a widened decision boundary, indicating pursuit of more evidence prior to choices. Our best-fitting model of DDM dynamics suggested that regardless of the state (low or high conflict), the boundary increased alongside increased goal-directed attention to both costs and rewards, as well as alongside increased cardiac contractility. However, neural and cardiac-sympathetic signals aligned to increasingly modulate boundary-width exclusively in states of high conflict, offering the first evidence of interactive cross-modal collaboration in value-based decisions. Analyses involving cross-trial coherence corroborated a conflict-specificity of neural-sympathetic relations and additionally proposed a role for sympathetics in disrupting the prepotent signal in reward-sensitivity.

Our findings suggest cardiac-sympathetic activity may therefore form a coordinated fireteam with neural processes to combat reward-sensitivity during the approach-avoidance conflict. Beginning with cardiac-sympathetics, the contractility-boundary relations are broadly consistent with sympathetic reactivity in contexts of increasing uncertainty^43^ and greater difficulty^2^. However, our cross-trial coherence findings are the strongest evidence yet that sympathetic reactivity might influence prepotent signal processing during value-based conflict. Across decision-making contexts, humans are usually biased toward more desirable information^44^, to the extent that an insensitivity to reward has been reported as a robust computational phenotype of psychiatric conditions such as depression^11,45^. In the present study, and in at least two separately reported human studies using the same task settings^9,10^, participants consistently over-weighted reward when making choices. More recent evidence using transiently disruptive cortical stimulation further proposes that reward-sensitivity might not simply reflect impulsivity, but a cortically-mediated model of a person’s primary goal in a value-based setting (i.e., capture reward)^12^. Under this assumption, it would be physiologically efficient to therefore prioritize reward information, and reserve effortful scrutiny and juxtaposition involving multiple streams of information for moments of conflict. Reward-sensitivity also generalizes to dynamic learning tasks, where recent studies report that people learn faster from positive-vs-negative prediction errors^5,46–47^. Consistent with our present findings, this learning asymmetry attenuates (i.e., learning from negative outcomes occurs more rapidly) when sympathetic activity is elevated^5,48^, to the extent that sympathetic reactivity even predicts individual participants who adjust their behavior more optimally to declining changes in their environment^5^. Whether cardiac-sympathetics serve common mechanisms to resolve uncertainty and address biases across decisions and learning is an exciting avenue of future research.

We additionally observed a collaborative contribution from neural dynamics in the alpha band. Broadly considered to reflect inhibition^49^ and visual spatial attention^50^, alpha power also shows a correspondingly flexible and goal-directed profile in cognitive processing. For example, during spatial recall, alpha power can code spatial targets in the absence of external information^51^ consistent with post-perceptual goal maintenance. If participants are cued to switch recall to a different memory location after memory arrays disappear, alpha dynamics can likewise switch from encoding the initial target to encoding the new one^52^. Alpha power can additionally signal a person’s willingness to take future risks^53^, suggesting it also responds in more value-based settings. Together, these findings are consistent with our interpretation that late alpha power mediated “fair assessment”, i.e., a shift in attention to process additional (cost) information alongside the prepotent signal (reward). Interestingly, we observed less association between steady-state visually-evoked potentials (SS) and DDM parameters. This might be due to task requirements. Earlier work implicates SS in coding information relevant for DDM decision boundaries^26^, albeit in tasks requiring perceptual and not value-based decisions. Our task used large visually unambiguous stimuli and created conflict that was value-based (subjective) rather than perceptually-driven. Recent human^54^ and nonhuman^55^ work dissociates alpha signals from modulating gain of sensory information, consistent with the idea that these signals have greater relevance for behavioral responses in value-based settings.

We lastly speculate on a network of substrates that might underly the behaviorally relevant interaction between the neural (alpha) and cardiac-sympathetic (contractility) signals in states of high conflict. It’s highly likely that our observed neural dynamics in the alpha band were facilitated by noradrenergic (NE) projections from the locus coeruleus in the brainstem (LC)^56–58^. The LC-NE system innervates cortical areas involved in orienting attention (e.g., parietal^59^), responding to arousal, goal-relevant stimuli and exploration^58^, all of which are likely relevant during moments of conflict. LC-NE can also broadly influence sympathetic activity^60^, however when it comes to specifically cardiac activity, evidence from both animal-optogenetic^61^ and human-imaging^62^ studies suggest LC-NE influences heart rate via vagal (i.e., parasympathetic) channels. In contrast, our specific cardiac assay - contractility (inotropy) - primarily tracks beta-adrenergic sympathetic drive to the heart when recorded from non-medicated physically stationary human participants (see discussion in^1^; also see methods for how, in our study, we corrected for influences of heart rate and respiratory cycle). A key subcortical controller of this cardiac-sympathetic response is the rostral ventrolateral medulla in the brainstem (RVLM)^63–65^, which is the primary source of organ-specific sympathetic preganglionic neurons. RVLM principally receives inputs from the cortically-modulated hypothalamus^67–69^. LC has few direct efferent connections with RVLM^66^, although it might communicate indirectly via its projections to the paraventricular nucleus of the hypothalamus^66^. The behavioral facilitation we observed when neural (alpha) and cardiac-sympathetic (contractility) signals interact may therefore reflect a nonlinear behavioral facilitation of two subcortical nodes (LC-NE and RVLM) activating concurrently. Alternatively, neural-cardiac-sympathetic collaboration may ultimately be mediated at the cortical level. Future inquiry could explore these alternatives with a view to first confirming involvement by LC-NE and RVLM in managing conflict, thereafter clarifying the specific roles performed by each, and finally describing the broader network structure of the response, i.e., the relevant functional afferents and efferents of each node.

### Concluding remarks

We reveal evidence that cardiac-sympathetic signals collaborate with neural processes to support humans making complex value-based decisions. We additionally reveal a potential mechanistic role for cardiac-sympathetic activity in disrupting processing of prepotent (reward) signals. Future research can explore whether cardiac-sympathetic responses serve common mechanisms across different cognitive tasks. The network of substrates underlying neural and cardiac-sympathetic collaboration in complex cognitive tasks also needs to be characterized. In terms of clinical relevance, autonomic function is vulnerable to neurodegenerative conditions such as Alzheimer’s and Parkinson’s disease^60,72^. Future research may therefore test if features of cross-modal collaboration during complex cognition can assist with early detection.

## Acknowledgements

Acknowledgments: The research was supported by award W911NF-16-1-0474 from the Army Research Office and by the Institute for Collaborative Biotechnologies under Cooperative Agreement W911NF-19-2-0026 with the Army Research Office.

## Author contributions

Conceptualization, N.M.D. and S.T.G.; Methodology, N.M.D., S.T.G. and J.O.G.; Software, N.M.D. and T.B.; Validation, N.M.D., T.B. and J.O.G.; Formal Analysis, N.M.D.; Investigation, A.S.; Resources, A.S., E.R. V.B. and D.Y.; Supervision, S.T.G., J.O.G., B.G., E.R. and V.B.; Writing - Original Draft, N.M.D.; Writing - Review & Editing, N.M.D., A.S., T.B., J.O.G., V.B., E.R., D.Y., B.G. and S.T.G.; Project Administration, S.T.G. and B.G.; Funding Acquisition, S.T.G. and B.G.

## STAR Methods

### KEY RESOURCES TABLE

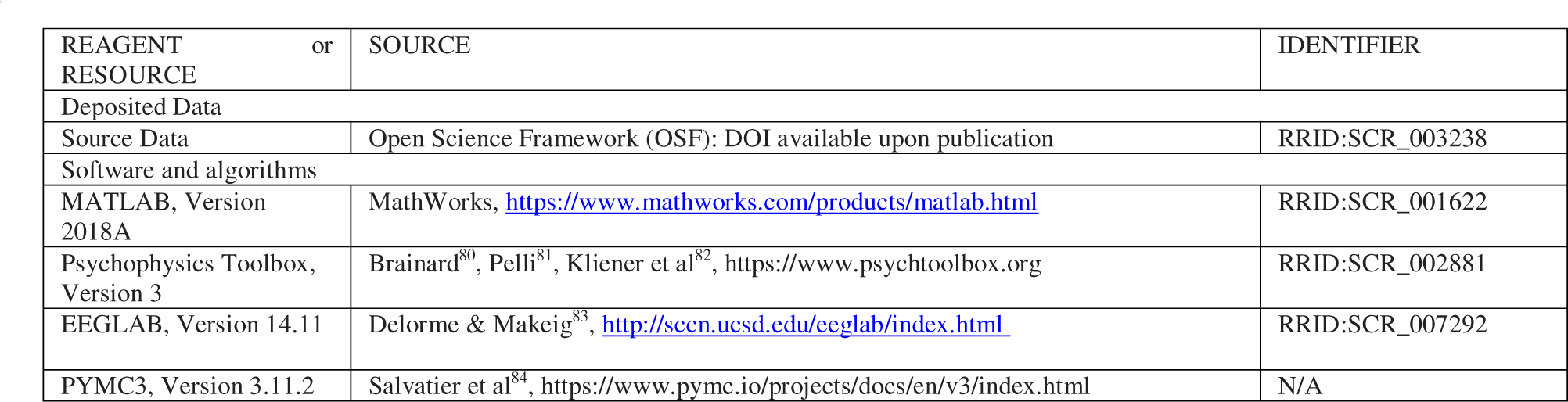
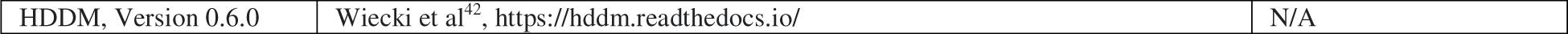

### RESOURCE AVAILABILITY

#### Lead contact

Further information and requests for data and code should be directed to and will be fulfilled by the lead contact, Neil M. Dundon.

#### Materials availability

This study did not generate new unique materials

#### Data and code availability

Data used in the reported analyses has been deposited at the Open Science Foundation (OSF) and is publicly available as of the date of publication. DOI is listed in the key resources table. Any additional information required to reanalyze the data reported in this paper is available from the lead contact upon request.

### EXPERIMENTAL MODEL AND STUDY PARTICIPANT DETAILS

We recruited an initial sample of 33 human participants, via both word-of-mouth and via an online participant recruitment portal operated by University of California, Santa Barbara (UCSB). We removed six participants from all analyses: one subject accepted more than 90% of offers, two participants’ EEG data had an artefact in more than 50% of epochs and one further subject satisfied both screening criteria. In addition, we removed two participants due to excessive noise in their impedance cardiography data. We accordingly report findings from a final sample of 27 participants. This group had a mean (standard deviation) age of 21.4 (3.3), and 17 were female. All participants were right-handed. Subject remuneration was $20 per hour base rate, with a bonus payment determined by their approach-avoidance behavior, which approximately corresponded to an additional $13.50 per subject. All testing took place during a single session in a quiet, dimly lit experimental suite and all procedures received approval from the Institutional Review Board at UCSB. Participants provided informed written consent, prior to participating.

### METHOD DETAILS

#### Approach-avoidance paradigm

We using a graded extension of the approach-avoidance framework previously employed in nonhuman primate^73,74^ and human^4,9–10^ contexts. Participants approach or avoid varying levels of monetary reward paired with varying levels of painful electric shock, in trial-by-trial “take-both- or-leave-both” offers.

Paradigm schematic is in Figure 2A. Participants made a total of 352 approach-avoidance choices (split into eight blocks of 44). Their head position was fixed by an adjustable chin and forehead rest, to maintain a viewing distance of 57 cm from the stimulus presentation screen: an ASUS VS278 monitor, viewing area 60 cm width by 33.5 cm height, refresh rate of 240 Hz (inter-frame interval=.004 s). We advised participants to move their bodies as little as possible, to prevent motion-related confounds entering the physiology recordings.

During each approach-avoidance trial described below, participants further fixated their eyes on a central point (RGB_[min=0,max=1]_=[.750 .750 .750]; diameter=0.221 °). The background color remained black (RGB_[min=0,max=1]_=[0 0 0]) at all times, except for payout trials (see below). Offers comprised four sequential events: (i) baseline, (ii) offer onset, (iii) offer offset and (iv) feedback. (i) Each offer initiated with a baseline period with a duration between 420 and 540 frames (inclusive) drawn with discrete uniform probability on each trial (approx. 1.75 s to 2.25 s). Baseline onset was signified by the immediate appearance of two vertically-oriented rectangular dot arrays, each spanning 7.30 ° width by 27.8 ° height, comprised of 79 columns and 322 rows of dots (dot diameter=.056°), with centroids positioned at a horizontal eccentricity +/- 3.75 ° from the central fixation point. (ii) Following baseline, during offer onset, the offer bars gradually communicated the magnitude of the offer’s value dimensions, with one bar communicating the level of offered reward and the other bar communicating the level of incurred shock. We drew a different offer on each trial from a two-dimensional decision (reward-by-shock) space, and communicated the magnitude of each dimension by gradually filling in an area of both bars with a relevant offer color (khaki or blue; one color per offer bar). Specifically, contiguous rows of dots, equally portioned above and below the centroid of each offer bar, gradually changed into one of two offer colors. The number of rows changing into an offer color indicated the magnitude offered in that dimension, i.e., the offered reward (no rows: $0.01, to all rows: $1.50) and the offered shock (no rows: minimum pain, to all rows: max bearable pain - see thresholding section below). Counterbalanced across participants, reward and shock mapped onto one color for the entire experiment, while color laterality was determined with 0.50 uniform probability before each trial. Offer onset duration was four seconds, with color change controlled by reward and shock gradients that respectively changed dot colors from baseline grey (RGB_[min=0,max=1]_=[.375 .375 .375]) to peak offer color (khaki, RGB_[min=0,max=1]_=[.632 .586 .351]; blue, RGB_[min=0,max=1]_=[.328 .616 .375]) at even step sizes in RGB space over the offer onset frames (n=960). (iii) Following peak onset, during offer offset, offer colors gradually returned to baseline grey over a two-second period. We instructed participants to respond only once as soon as they decided whether to approach or avoid an offer, with a time limit of the end of offer offset. Participants registered their responses by pressing z with the index finger of their left hand, or m with the index finger of their right hand. The mapping of z and m onto approach and avoidance responses was fixed for each block of 44 trials, determined prior to the block with 0.50 uniform probability. (iv) Following offer offset, participants observed a feedback screen for one second (nine seconds for payout trials, see below). As depicted in the lower portion of Figure 2A, a “successful” response, i.e., a single response executed during offer onset or offset, led to a confirmation feedback screen containing a snapshot image of the offer (at peak offer colors), accompanied by a written confirmation of their choice. If participants responded more than once, responded during the baseline (pre-onset), or failed to execute any response before the final frame of offer offset, a warning appeared on the screen indicating the relevant error, along with the lateralized response prompts (lower portion of Figure 2A). We stored all error trials and reissued them to participants after the eighth block, to ensure that errors could not be a strategy to circumvent specific offers. All feedback screens (successful, error or payout (below)) also included prompts to remind participants which colors mapped onto the different reward dimensions (indicated with a lightning bolt (shock) or dollar symbol (reward) overlaying the relevant bar), and which button response mapped onto which choice for the given block. This screen was also displayed prior to each block. Finally, offer bars flickered throughout each trial, either left: 12Hz, right: 13.33Hz or vice versa, determined with 0.50 uniform probability. We presented all stimuli with customized scripts in MATLAB (Version 2018a, The Mathworks Inc., Natick, MA, USA, https://www.mathworks.com/products/matlab.html) using functions from PsychToolbox-3^80–82^. Successfully registered choices were coded either 1 (approach) or 0 (avoid), and response time (RT) was the time (in seconds) between offer onset and choice execution.

#### Payout trials

For safety reasons, we did not administer any electric shocks during the testing session while participants wore physiology electrodes. We instead postponed payout on an infrequent subset of trials, in line with previous work recording physiology signals during similar approach-avoidance paradigms^4,9^. Prior to testing, we selected eighteen “payout trials” (5.11%) with uniform probability across the entire set of offers provided the offered shock was below 80% of a subject’s maximum pain level. The baseline, onset and offset sequence of payout trials were identical to non-payout trials, however, during the feedback section of payout trials (extended from one to nine seconds), the screen changed from black to red (RGB_[min=0,max=1]_=[1 0 0]; left lower portion of Fig. 1c) and participants learned that the monetary reward and shock from that offer would be administered following the testing session (were that offer approached) or that the values would have been administered (were that offer avoided). If participants made a response error during a payout trial, they instead saw the error feedback screen, and the payout trial was added to the list of trials to be reissued. We instructed participants that payout trials could not be predicted before registering a response and to treat each offer as a potential payout trial. Previous work^4,9^ reports that payout trials do not affect behavior on subsequent choices.

#### Neural recordings

Concurrent with the approach-avoidance paradigm, we recorded continuous electroencephalogram (EEG) data from a montage of 63 scalp electrodes (channels) arranged using the International 10-20 system. We sampled the EEG signal at 1000 Hz from each channel, using a BrainAmp MR amplifier (Brain Products, Berlin, Germany). Channel FCz served as the online reference while channel Cz served as the ground. Between blocks, experimenters paused recordings to check electrode impedance and noisy channels.

#### Cardiac-sympathetic recordings

Concurrent with the approach-avoidance paradigm we also recorded data from combined electrocardiogram (EKG) and impedance cardiogram (ICG) using a total of ten EL500 electrodes (BIOPAC, USA). All electrode sites were prepped with an exfoliating pad and NuPrep gel. We placed one electrode beneath the right collarbone and one beneath the left rib cage to record EKG. We placed eight electrodes to both record ICG and serve as the ground: two on each side of the torso and two on each side of the neck. We sampled both the EKG and ICG at 5000Hz respectively using ECG100C and NICO100C amplifiers, integrated with an MP150 system (BIOPAC, USA). Online, AcqKnowledge software version 4.3 differentiated the raw (z) ICG data with respect to time (dz/dt) and removed respiratory artefact from the ensuing dz/dt waveform with a high-pass filter (BIOPAC, USA). Between blocks, experimenters paused recordings to check for noise in the EKG and ICG data.

#### EEG preprocessing

EEG preprocessing used functions available in the EEGLAB toolbox^83^. First, each participant’s EEG data were downsampled (250Hz) and hi-pass filtered (<1Hz) separately for each block. Line noise was removed with an automated function^76^. We merged resulting sets of blockwise data into a single set (one set per subject) and identified noisy channels using an automated function that tested whether data in each channel correlated with those in surrounding channels by a coefficient of at least 0.85^77^. Identified channels were replaced using spherical interpolation. We then re-referenced datasets to the montage average, created epochs spanning from -1 s to +6 s relative to offer onset, and subtracted baseline means (taken from the window -.5 s to 0 s relative to offer onset). We then performed ICA decomposition separately on each subject’s resulting epochs, and stored the resulting weights of components that were 0.95 likely to be ocular or cardiac activity, determined by an automated classifier^78^. We next imported, downsampled, hi-pass filtered and removed line noise from participants’ separate blocks of raw data again, as above. Separately for each block, we replaced noisy channels as above and removed ICA components related to ocular and cardiac artefacts. We marked any data point where any channel still exceeded 150 mV (for later rejection) and applied a spatial Laplacian filter across multichannel data at each time point. We then reversed the laterality of electrodes on all trials where reward appeared on the left of the screen, so that each trial “de facto” presented reward on the right and cost on the left. We refer to data at this stage as “preprocessed” data.

To extract timeseries from preprocessed data for steady-state visually evoked potentials relevant for reward (SS_rew_) and cost (SS_shk_) information (Figure 3A), we used rectified and smoothed power timeseries that had been filtered to either 12 or 13.33 Hz, depending on the flicker of reward or cost information for a given trial (note that no specific frequency mapped onto either reward or cost; flickers varied trial-by-trial). We convolved each channel’s fast-Fourier transformed data with trapezoid-shaped bandpass filters (“on” width = .5 Hz, transition bandwidth = 0.25 Hz), centered on 12 Hz or 13.33 Hz before rectifying, smoothing (mean within sliding windows spanning 66 ms) and downsampling inverse-fourier timeseries to 125 Hz. We also constructed a third dataset, using these exact procedures, but with filters cantered on 12.66 Hz (midway between 12 and 13.33 Hz), and subtracted it from SS_rew_ and cost SS_shk_ timeseries to mitigate SS influence from underlying activity in the alpha band. We created epochs spanning from -1 s to +6 s relative to offer onset for the 12 and 13.33 Hz datasets, removing the baseline average value in the 500 ms window prior to offer onset. For trial-by-trial measures, we computed the average power in early ([0 s to 1 s] post offer onset) and late ([1 s to 2 s]) time windows (Figure 3A). Amplitudes were averaged at electrode sites contralateral to the relevant information, i.e., O1 and PO1, or at electrodes O2 and PO2, depending on reward and cost laterality on a given trial (Figure 3A).

We also extracted timeseries from preprocessed data of spatially-sensitive alpha power, i.e., relevant for reward (alpha_rew_) and cost (alpha_shk_) information (Figure 3B). We first applied notch filters to remove power in SS frequencies (inverse of bandpass filters above), and then convolved each channel’s fast-Fourier transformed data with a trapezoid-shaped bandpass filter (“on” segment spanning 7 Hz to 14 Hz, transition bandwidth = 0.5 Hz). Resulting inverse-fourier timeseries were rectified, smoothed (mean within sliding windows spanning 66 ms) and downsampled channels to 125 Hz. We created epochs spanning from -1 s to +6 s relative to offer onset, removing the baseline average value in the 500 ms window prior to offer onset. For trial-by-trial measures, we computed the average power in early ([0 s to 1 s] post offer onset) and late ([1 s to 2 s]) time windows (Figure 3B). Amplitudes were averaged at electrode sites contralateral to the relevant information, i.e., PO and PO7, or at electrodes PO2 and PO8, depending on reward and cost laterality on a given trial (Figure 3B).

For both the SS and alpha timeseries we also computed symmetry timeseries at each timepoint (t); i..e, SS(t)_sym_=-1*log(|SS(t)_rew_-SS(t)_rew_|); alpha(t)_sym_=-1*log|alpha(t)_rew_-alpha(t)_shk_|). For trial-by-trial measures, we computed averages in the same manner as the single traces. Higher values reflecting higher symmetry.

We computed alpha-phase timeseries from preprocessed data using the same steps above for the power timeseries, up to the convolution of the bandpass filter on each fast-Fourier transformed channel. From there, we computed the phase angle at each time point using the angle function in MATLAB. Phase timeseries (alpha_rew_θ and alpha_shk_θ) were extracted at electrode sites contralateral to the relevant information, i.e., PO and PO7, or at electrodes PO2 and PO8, depending on reward and cost laterality on a given trial.

#### EKG/ICG preprocessing and contractility assay

We estimated contractility from the pre-ejection period (PEP). Using a semi-automated software package (MEAP^75^), we identified the R point of the EKG QRS complex (early systole: initial left-ventricular depolarization) and the B point of the dz/dt waveform from the ICG (mid systole: opening of the aortic valve), for each individual heartbeat (Figure 3C). The time-period between these two cardiac events is the pre-ejection. This electro-mechanical time interval, covering systolic activity from the initial electrical depolarization of the left ventricle until the opening of the aortic valve, is an index of beta-adrenergic contraction vigor, and is primarily mediated by sympathetic activity^35–40^. Shorter intervals reflect increased contractility (positive inotropy). For trial-by-trial measures, we computed the average PEP value across all heartbeats occurring in the two-second window immediately following offer onset (Figure 3C). These values were log-transformed and then reverse-scored, so that higher values reflect higher contractility.

### QUANTIFICATION AND STATISTICAL ANALYSIS

#### Estimating conflict

We fitted a separate logistic model to each individual subject’s trial-by-trial data, modeling the probability of approaching each offer (t) as: s’(SV), where s’(x) is sigmoid transition function: 1/(1+e^-x^), and SV=(β_0_+β_1_*reward_t_+β_2_*shock_t_). We fitted each model using the mnrfit function in MATLAB, after first normalizing reward and shock to z-score ranges within participants. In this way SV>0 reflects p(approach)>0.50 and vice-versa. From trial-by-trial SV estimates we could either compute a continuous measure of conflict, i.e., the degree of equivalence in the value of approaching or avoiding each offer, with function c’(SV), where c’(x) = -1*z’(log(x^2^)) and z’(x) is a z-score. We also measured a discrete conflict “state”. For this, we separately median split each participant’s approach and avoid trials by SV and classified low conflict (approach above median SV, avoid below median SV) and high conflict (approach below median SV, avoid above median SV). We estimated SV, continuous conflict and conflict states for each trial prior to any screening due to EEG or sympathetic artefact.

#### Modeling trial-by-trial RT

To find the best fitting model of participants’ log-transformed trial-by-trial RT, we we ran a series of Bayesian linear mixed-effects regression models, combining (4Ck, for k∈{1,…,4}) the following four coefficients: reward (β_1_), cost (β_2_), subjective value (SV;β_3_) and conflict (AAC; β_4_). Model fits were assessed with the widely accepted inference criterion (WAIC). From the best fitting model, we report group-level coefficient posteriors. We fitted these models to trials that were screened for EEG or sympathetic artefact. Posteriors were sampled and WAICs were computed using pymc3 functions^84^ in Python.

#### Modeling choice inconsistency

Consistent approach behavior is to approach positive SV (p(approach)>0.50) and consistent avoid behavior is to avoid negative SV (p(approach)<0.50). We compared the proportion of subjects’ consistent choices between states of high and low conflict (separately for approach and avoid choices) at the group level using a paired-samples t-test.

#### Computational modeling

We fitted hierarchical Bayesian drift-diffusion models (DDM; Figure 1D) to choice and RT data to estimate group level parameters for boundary separation (a), drift rate (v), starting point (z) and nondecision time (t). Discrete and trial-by-trial modulation of parameters was administered using a regression procedure^42^. The lower boundary was avoid and the higher boundary was approach. A single drift rate was fitted using a link function that made it negative on avoid trials and positive on approach trials.

In the baseline model reported in Figure 2E, we fitted discrete parameters for trials in states of high and low conflict.

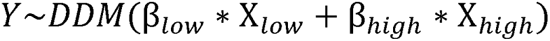

Where Y is the vector of trial-by-trial choice and RT data, each β*_low_* and β*_high_* each contain four posterior estimates, i.e., of DDM parameters [a,v,z,t]. X*_low_* and X*_high_* are dummy-coded design vectors respectively for states of low and high conflict and DDM(x) is the Weiner diffusion likelihood function described in^79^.

In models reported in Figure 3D-F, we also included coefficient estimates that tested if the parameters in β*_low_* and β_1_ varied as a function of trial-by-trial physiological signals:

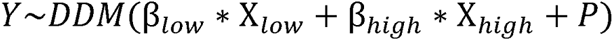

Where Y, β*_low_*, β*_high_*, X*_low_*, X*_high_* and DDM(x) are as above, and P contained:

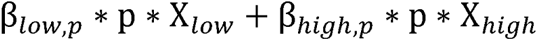

for each signal (p) included in the model. β*_low,p_* and β*_high,p_* each contain four coefficient posterior estimates, i.e., describing the relation between DDM parameters [a,v,z,t] and signal (p) on each trial.

In each model, we sampled group-level posteriors for β*_low_*, β*_high_*, β*_low,p_* and β*_high,p_* 5000 times with Markov-Chain Monte Carlo, using the HDDMRegressor function from the HDDM toolbox^42^ version 0.6.0 in python 2.7. We discarded the first 500 samples of each posterior estimate as tuning steps. Model fits were assessed with Deviance Inference Scores (DIC).

#### Alpha-phase coherence

We first averaged across trials for each participant’s alpha_rew_θ and alpha_shk_θ timeseries, separately for trials in states of high and low conflict and separately again for trials in states of high and low contractility (determined by a median split across all of a participants trials). To compute a summary estimate of early and late coherence across trials, we rectified the resulting participant-average phase waveforms |θ|, and computed the average values across datapoints in early ([0 s to 1 s] post offer onset) and late ([1 s to 2 s]) time windows (Figure 4A). For trials in states of low and high conflict we separately tested the effects of timeseries (alpha_rew_θ, alpha_shk_θ), time window (early, late) and contractility (high, low) across summarized rectified phase angles with three-way within-subjects ANOVAs.

## Notes

### Competing Interest Statement

The authors have declared no competing interest.

## References

1. Stump, A., Gregory, C., Babenko, V., Rizor, E., Bullock, T., Macy, A., Giesbrecht, B., Grafton, S. T., & Dundon, N. M. (2023). Non-invasive monitoring of cardiac contractility: Trans-radial electrical bioimpedance velocimetry (TREV). Psychophysiology, 00, e14411.

2. Richter, M., Friedrich, A., & Gendolla, G. H. (2008). Task difficulty effects on cardiac activity. Psychophysiology, 45(5), 869–875.

3. Richter, M., Gendolla, G. H. E., & Wright, R. A. (2016). Three decades of research on motivational intensity theory: What we have learned about effort and what we still don’t know. Advances in motivation science, 3, 149–186.

4. Dundon, N. M., Shapiro, A. D., Babenko, V., Okafor, G. N., & Grafton, S. T. (2021). Ventromedial prefrontal cortex activity and sympathetic allostasis during value-based ambivalence. Frontiers in Behavioral Neuroscience, 15, 615796.

5. Dundon, N. M., Garrett, N., Babenko, V., Cieslak, M., Daw, N. D., & Grafton, S. T. (2020). Sympathetic involvement in time-constrained sequential foraging. *Cognitive, Affective*, & Behavioral Neuroscience, 20, 730–745.

6. Champion, R. A. (1961). Motivational effects in approach-avoidance conflict. Psychological Review, 68(5), 354.

7. Elliot, A. J., & Thrash, T. M. (2002). Approach-avoidance motivation in personality: approach and avoidance temperaments and goals. Journal of Personality and Social Psychology, 82(5), 804.

8. Zorowitz, S., Rockhill, A. P., Ellard, K. K., Link, K. E., Herrington, T., Pizzagalli, D. A., Widge, A. S., Deckersbach, T., & Dougherty, D. D. (2019). The neural basis of approach-avoidance conflict: A model based analysis. Eneuro, 6(4).

9. Shapiro, A. D., & Grafton, S. T. (2020). Subjective value then confidence in human ventromedial prefrontal cortex. Plos One, 15(2), e0225617.

10. Volz, L. J., Welborn, B. L., Gobel, M. S., Gazzaniga, M. S., & Grafton, S. T. (2017). Harm to self outweighs benefit to others in moral decision making. Proceedings of the National Academy of Sciences, 114(30), 7963–7968.

11. Pedersen, M. L., Ironside, M., Amemori, K. I., McGrath, C. L., Kang, M. S., Graybiel, A. M., & Frank, M. J. (2021). Computational phenotyping of brain-behavior dynamics underlying approach-avoidance conflict in major depressive disorder. PLoS computational biology, 17(5), e1008955.

12. Rolle, C. E., Pedersen, M. L., Johnson, N., Amemori, K. I., Ironside, M., Graybiel, A. M., & Etkin, A. (2022). The role of the dorsal–lateral prefrontal cortex in reward sensitivity during approach–avoidance conflict. Cerebral Cortex, 32(6), 1269–1285.

13. Talmi, D., Dayan, P., Kiebel, S. J., Frith, C. D., & Dolan, R. J. (2009). How humans integrate the prospects of pain and reward during choice. Journal of Neuroscience, 29(46), 14617–14626.

14. Park, S. Q., Kahnt, T., Rieskamp, J., & Heekeren, H. R. (2011). Neurobiology of value integration: When value impacts valuation. Journal of Neuroscience, 31(25), 9307–9314.

15. Skvortsova, V., Palminteri, S., & Pessiglione, M. (2014). Learning to minimize efforts versus maximizing rewards: Computational principles and neural correlates. Journal of Neuroscience, 34(47), 15621–15630.

16. Wan, X., Cheng, K., & Tanaka, K. (2015). Neural encoding of opposing strategy values in anterior and posterior cingulate cortex. Nature Neuroscience, 18(5), 752–759.

17. Usher, M., & McClelland, J. L. (2001). The time course of perceptual choice: the leaky, competing accumulator model. Psychological review, 108(3), 550.

18. Ratcliff, R., & McKoon, G. (2008). The diffusion decision model: theory and data for two-choice decision tasks. Neural computation, 20(4), 873–922.

19. Forstmann, B. U., Ratcliff, R., & Wagenmakers, E. J. (2016). Sequential sampling models in cognitive neuroscience: Advantages, applications, and extensions. Annual review of psychology, 67, 641–666.

20. Peters, J., & D’Esposito, M. (2020). The drift diffusion model as the choice rule in inter-temporal and risky choice: A case study in medial orbitofrontal cortex lesion patients and controls. PLoS computational biology, 16(4), e1007615.

21. Shahar, N., Hauser, T. U., Moutoussis, M., Moran, R., Keramati, M., Nspn Consortium, & Dolan, R. J. (2019). Improving the reliability of model-based decision-making estimates in the two-stage decision task with reaction-times and drift-diffusion modeling. PLoS computational biology, 15(2), e1006803.

22. Ballard, I. C., & McClure, S. M. (2019). Joint modeling of reaction times and choice improves parameter identifiability in reinforcement learning models. Journal of Neuroscience Methods, 317, 37–44.

23. Colas, J. T. (2017). Value-based decision making via sequential sampling with hierarchical competition and attentional modulation. PloS one, 12(10), e0186822.

24. Fontanesi, L., Gluth, S., Spektor, M. S., & Rieskamp, J. (2019). A reinforcement learning diffusion decision model for value-based decisions. Psychonomic bulletin & review, 26(4), 1099–1121.

25. Dundon, N. M., Colas, J. T., Garrett, N., Babenko, V., Rizor, E., Yang, D., & Grafton, S. T. (2023). Decision heuristics in contexts integrating action selection and execution. Scientific Reports, 13(1), 6486.

26. O’connell, R. G., Dockree, P. M., & Kelly, S. P. (2012). A supramodal accumulation-to-bound signal that determines perceptual decisions in humans. Nature neuroscience, 15(12), 1729–1735.

27. Galloway, N. R. (1990). Human brain electrophysiology: Evoked potentials and evoked magnetic fields in science and medicine. The British Journal of Ophthalmology, 74(4), 255.

28. Müller, M. M., Picton, T. W., Valdes-Sosa, P., Riera, J., Teder-Sälejärvi, W. A., & Hillyard, S. A. (1998). Effects of spatial selective attention on the steady-state visual evoked potential in the 20–28 Hz range. Cognitive Brain Research, 6(4), 249–261.

29. Müller, M. M., Andersen, S., Trujillo, N. J., Valdes-Sosa, P., Malinowski, P., & Hillyard, S. (2006). Feature-selective attention enhances color signals in early visual areas of the human brain. Proceedings of the National Academy of Sciences, 103(38), 14250–14254.

30. Gulbinaite, R., Roozendaal, D. H., & VanRullen, R. (2019). Attention differentially modulates the amplitude of resonance frequencies in the visual cortex. NeuroImage, 203, 116146.

31. Pfurtscheller G, & Aranibar A. (1977). Event-related cortical desynchronization detected by power measurements of scalp EEG. Electroencephalography and Clinical Neurophysioly, 42(6), 817–26.

32. Foxe, J. J., & Snyder, A. C. (2011). The role of alpha-band brain oscillations as a sensory suppression mechanism during selective attention. Frontiers in Psychology, 2, 154.

33. Klimesch, W. (2012). Alpha-band oscillations, attention, and controlled access to stored information. Trends in Cognitive Sciences, 16(12), 606–617.

34. Wang, C., Rajagovindan, R., Han, S. M., & Ding, M. (2016). Top-down control of visual alpha oscillations: sources of control signals and their mechanisms of action. Frontiers in human neuroscience, 10, 15.

35. Lewis, R. P., Leighton, R. F., Forester, W. F., & Weissler, A. M. (1974). Systolic time intervals. In A. M. Weissler (Ed.), Noninvasive cardiology (pp. 301–368). Grune and Stratton.

36. Light, K. C. (1985). Cardiovascular and renal responses to competitive mental challenges. In J. F. Orlebeke, G. Mulder, & L. J. P. van Doornen (Eds.), Cardiovascular psychophysiology: Theory and methods (pp. 683–702). Plenum.

37. Linden, W. (1985). Cardiovascular response as a function of predisposition, coping behavior and stimulus type. Journal of Psychosomatic Research, 29(6), 611–620.

38. Newlin, D. B., & Levenson, R. W. (1979). PreLJejection period: Measuring betaLJadrenergic influences upon the heart. Psychophysiology, 16(6), 546–552.

39. Sherwood, A., Allen, M. T., Obrist, P. A., & Langer, A. W. (1986). Evaluation of betaLJadrenergic influences on cardiovascular and metabolic adjustments to physical and psychological stress. Psychophysiology, 23(1), 89–104.

40. Sherwood, A., Allen, M. T., Fahrenberg, J., Kelsey, R. M., Lovallo, W. R., & Van Doornen, L. J. (1990). Methodological guidelines for impedance cardiography. Psychophysiology, 27(1), 1–23.

41. Callister, R., Suwarno, N. O., & Seals, D. R. (1992). Sympathetic activity is influenced by task difficulty and stress perception during mental challenge in humans. The Journal of Physiology, 454(1), 373–387.

42. Wiecki, T. V., Sofer, I., & Frank, M. J. (2013). HDDM: Hierarchical Bayesian estimation of the drift-diffusion model in Python. Frontiers in neuroinformatics, 7(*14*), 1–10.

43. Palacios-Filardo, J., & Mellor, J. R. (2019). Neuromodulation of hippocampal long-term synaptic plasticity. Current opinion in neurobiology, 54, 37–43.

44. Sharot, T., & Garrett, N. (2016). Forming beliefs: Why valence matters. Trends in Cognitive Sciences, 20(1), 25–33.

45. Garrett, N., Sharot, T., Faulkner, P., Korn, C. W., Roiser, J. P., & Dolan, R. J. (2014). Losing the rose tinted glasses: neural substrates of unbiased belief updating in depression. Frontiers in human neuroscience, 8, 639.

46. Garrett, N., & Daw, N. D. (2020). Biased belief updating and suboptimal choice in foraging decisions. Nature communications, 11(1), 3417.

47. Lefebvre, G., Lebreton, M., Meyniel, F., Bourgeois-Gironde, S., & Palminteri, S. (2017). Behavioural and neural characterization of optimistic reinforcement learning. Nature Human Behaviour, 1(4), 0067.

48. Garrett, N., González-Garzón, A. M., Foulkes, L., Levita, L., & Sharot, T. (2018). Updating beliefs under perceived threat. Journal of Neuroscience, 38(36), 7901–7911.

49. Jensen, O., & Mazaheri, A. (2010). Shaping functional architecture by oscillatory alpha activity: gating by inhibition. Frontiers in human neuroscience, 4, 186.

50. Worden, M. S., Foxe, J. J., Wang, N., & Simpson, G. V. (2000). Anticipatory biasing of visuospatial attention indexed by retinotopically specific alpha-band electroencephalography increases over occipital cortex. The Journal of neuroscience: the official journal of the Society for Neuroscience, 20(6), RC63–RC63.

51. MacLean, M. H., Bullock, T., & Giesbrecht, B. (2019). Dual process coding of recalled locations in human oscillatory brain activity. Journal of Neuroscience, 39(34), 6737–6750.

52. van Moorselaar, D., Foster, J. J., Sutterer, D. W., Theeuwes, J., Olivers, C. N., & Awh, E. (2018). Spatially selective alpha oscillations reveal moment-by-moment trade-offs between working memory and attention. Journal of cognitive neuroscience, 30(2), 256–266

53. Zhang, D., & Gu, R. (2018). Behavioral preference in sequential decision making and its association with anxiety. Human brain mapping, 39(6), 2482–2499.

54. Zhigalov, A., & Jensen, O. (2020). Alpha oscillations do not implement gain control in early visual cortex but rather gating in parietoLJoccipital regions. Human Brain Mapping, 41(18), 5176–5186.

55. Bastos, A. M., Lundqvist, M., Waite, A. S., Kopell, N., & Miller, E. K. (2020). Layer and rhythm specificity for predictive routing. Proceedings of the National Academy of Sciences, 117(49), 31459–31469.

56. Rajkowski, J. (1993). Correlations between locus coeruleus (LC) neural activity, pupil diameter and behavior in monkey support a role of LC in attention. Soc. Neurosc., Abstract, Washington, DC, 1993.

57. Joshi, S., & Gold, J. I. (2020). Pupil size as a window on neural substrates of cognition. Trends in cognitive sciences, 24(6), 466–480.

58. Aston-Jones, G., & Cohen, J. D. (2005). An integrative theory of locus coeruleus-norepinephrine function: adaptive gain and optimal performance. Annu. Rev. Neurosci., 28, 403–450.

59. Foote, S. L., & Morrison, J. H. (1987). Extrathalamic modulation of cortical function. Annual review of neuroscience, 10(1), 67–95.

60. Samuels, E. R., & Szabadi, E. R. S. A. E. (2008). Functional neuroanatomy of the noradrenergic locus coeruleus: its roles in the regulation of arousal and autonomic function part II: physiological and pharmacological manipulations and pathological alterations of locus coeruleus activity in humans. Current neuropharmacology, 6(3), 254–285.

61. Wang, X., Piñol, R. A., Byrne, P., & Mendelowitz, D. (2014). Optogenetic stimulation of locus ceruleus neurons augments inhibitory transmission to parasympathetic cardiac vagal neurons via activation of brainstem α1 and β1 receptors. Journal of Neuroscience, 34(18), 6182–6189.

62. Wood, C. S., Valentino, R. J., & Wood, S. K. (2017). Individual differences in the locus coeruleus-norepinephrine system: Relevance to stress-induced cardiovascular vulnerability. Physiology & behavior, 172, 40–48.

63. Shapoval, L. N., Sagach, V. F., & Pobegailo, L. S. (1991). Chemosensitive ventrolateral medulla in the cat: the fine structure and GABA-induced cardiovascular effects. Journal of the autonomic nervous system, 36(3), 159–172.

64. Mandal, A. K., Kellar, K. J., Norman, W. P., & Gillis, R. A. (1990). Stimulation of serotonin2 receptors in the ventrolateral medulla of the cat results in nonuniform increases in sympathetic outflow. Circulation research, 67(5), 1267–1280.

65. Kulkarni, S. S., Mischel, N. A., & Mueller, P. J. (2023). Revisiting differential control of sympathetic outflow by the rostral ventrolateral medulla. Frontiers in Physiology, 13, 2761.

66. Samuels, E. R., & Szabadi, E. (2008). Functional neuroanatomy of the noradrenergic locus coeruleus: its roles in the regulation of arousal and autonomic function part I: principles of functional organisation. Current neuropharmacology, 6(3), 235–253.

67. Kono, Y., Yokota, S., Fukushi, I., Arima, Y., Onimaru, H., Okazaki, S., & Okada, Y. (2020). Structural and functional connectivity from the dorsomedial hypothalamus to the ventral medulla as a chronological amplifier of sympathetic outflow. Scientific reports, 10(1), 13325.

68. Koba, S., Kumada, N., Narai, E., Kataoka, N., Nakamura, K., & Watanabe, T. (2022). A brainstem monosynaptic excitatory pathway that drives locomotor activities and sympathetic cardiovascular responses. Nature Communications, 13(1), 5079.

69. Dum, R. P., Levinthal, D. J., & Strick, P. L. (2019). The mind–body problem: Circuits that link the cerebral cortex to the adrenal medulla. Proceedings of the National Academy of Sciences, 116(52), 26321–26328.

70. Huangfu, D., Verberne, A. J., & Guyenet, P. G. (1992). Rostral ventrolateral medullary neurons projecting to locus coeruleus have cardiorespiratory inputs. Brain research, 598(1-2), 67–75.

71. Sara, S. J., & Bouret, S. (2012). Orienting and reorienting: the locus coeruleus mediates cognition through arousal. Neuron, 76(1), 130–141

72. Engelender, S., & Isacson, O. (2017). The threshold theory for Parkinson’s disease. Trends in neurosciences, 40(1), 4–14.

73. Amemori, K. I., & Graybiel, A. M. (2012). Localized microstimulation of primate pregenual cingulate cortex induces negative decision-making. Nature Neuroscience, 15(5), 776–785.

74. Amemori, K. I., Amemori, S., & Graybiel, A. M. (2015). Motivation and affective judgments differentially recruit neurons in the primate dorsolateral prefrontal and anterior cingulate cortex. Journal of Neuroscience, 35(5), 1939–1953.

75. Cieslak, M., Ryan, W. S., Babenko, V., Erro, H., Rathbun, Z. M., Meiring, W., & Grafton, S. T. (2018). Quantifying rapid changes in cardiovascular state with a moving ensemble average. Psychophysiology, 55(4), e13018.

76. Mullen T. (2012). NITRC: CleanLine: Tool/Resource Info. http://www.nitrc.org/projects/cleanline

77. Kothe, C. A., & Makeig, S. (2013). BCILAB: A platform for brain–computer interface development. Journal of Neural Engineering, 10(5), 056014.

78. Pion-Tonachini, L., Kreutz-Delgado, K., & Makeig, S. (2019). ICLabel: An automated electroencephalographic independent component classifier, dataset, and website. NeuroImage, 198, 181–197.

79. Navarro, D. J., & Fuss, I. G. (2009). Fast and accurate calculations for first-passage times in Wiener diffusion models. Journal of mathematical psychology, 53(4), 222–230.

80. Brainard, D. H. (1997) The Psychophysics Toolbox, Spatial Vision 10:433–436.

81. Pelli, D. G. (1997) The VideoToolbox software for visual psychophysics: Transforming numbers into movies, Spatial Vision 10:437–442.

82. Kleiner M, Brainard D, Pelli D, 2007, “What’s new in Psychtoolbox-3?” Perception 36 ECVP Abstract Supplement.

83. Delorme, A., & Makeig, S. (2004). EEGLAB: an open source toolbox for analysis of single-trial EEG dynamics including independent component analysis. Journal of neuroscience methods, 134(1), 9–21.

84. Salvatier, J., Wiecki, T. V., & Fonnesbeck, C. (2016). Probabilistic programming in Python using PyMC3. PeerJ Computer Science, 2, e55.

